# *In vivo* magnetic recording of single-neuron action potentials

**DOI:** 10.1101/2023.06.30.547194

**Authors:** Frederike J. Klein, Patrick Jendritza, Chloé Chopin, Mohsen Parto-Dezfouli, Aurélie Solignac, Claude Fermon, Myriam Pannetier-Lecoeur, Pascal Fries

## Abstract

Measuring fast neuronal signals is the domain of electrophysiology and magnetophysiology. While electrophysiology is easier to perform, magnetophysiology avoids tissue-based distortions and measures a signal with directional information. At the macroscale, magnetoencephalography (MEG) is established, and at the mesoscale, visually evoked magnetic fields have been reported. At the microscale however, while benefits of recording magnetic counterparts of electric spikes would be numerous, they are also highly challenging *in vivo*. Here, we combine magnetic and electric recordings of neuronal action potentials in anesthetized rats using miniaturized giant magneto-resistance (GMR) sensors. We reveal the magnetic signature of action potentials of well-isolated single units. The recorded magnetic signals showed a distinct waveform and considerable signal strength. This demonstration of *in vivo* magnetic action potentials opens a wide field of possibilities to profit from the combined power of magnetic and electric recordings and thus to significantly advance the understanding of neuronal circuits.

**Significance statement:** Electrophysiological tools allow the measurement of single-neuron action potentials with high temporal resolution. Magnetophysiological measurements would add valuable information, but are particularly hard to achieve for single neurons. Established technology for non-invasive magnetic brain signal measurements can currently not be used inside living tissue. Here, we demonstrate that miniaturized magnetic sensors based on giant magneto-resistance enable the measurement of the magnetic counterpart of single-neuron action potentials in vivo. This proof-of-principle shows a way towards integrating magnetic and electric recordings in future experiments and thus to profit from the complementary information measured by the two modalities.

## Introduction

Neuroscience is often driven by the development of new methods to record neuronal activity. Neuronal activity is characterized by electric currents, generated by neuronal outputs in the form of action potentials, and by neuronal inputs in the form of synaptic potentials. Synaptic potentials are often highly correlated across thousands to millions of neighboring synapses, thereby summing effectively to mass potentials, which are recorded as LFP (and LFP-derived signals like ECoG and EEG). By contrast, action potentials are typically weakly correlated between neighboring neurons, and are often studied using extracellular microelectrode recordings of so called “spikes”, which can provide information on the level of single neurons or small clusters of neurons. Spike recordings provide unique information about neuronal outputs, often revealing high spatial specificity.

Electric currents are inextricably linked to magnetic fields (Fig 1A). Thus, magnetic equivalents must exist both for action potentials and LFPs. However, *in vivo*, only LFP equivalents have been magnetically recorded, in the form of MEG. MEG sensors need to be extremely sensitive (in the femto-Tesla range) and massively shielded, to detect the weak neuronally generated fields. The development of this technology was highly challenging (Cohen, 1968; Hari et al., 1984; Hämäläinen et al., 1993; Hari and Salmelin, 1997), and still today, MEG is technologically much more demanding and expensive than its electric counterpart EEG. Nevertheless, MEG is used at many research institutions, because of its specific advantages: (1) While electrical currents need to flow through tissue, thereby getting attenuated, distorted and smeared by the tissue’s variable conductivity, magnetic signals pass through the neuropil almost unaffected (Barnes and Greenebaum, 2007) (2) While electrical recordings in practice require a reference, magnetic recordings do not. Reference-free recordings are particularly advantageous for analyses of functional connectivity based on correlations, which can be spuriously introduced by shared references. (3) While electrical recordings merely provide a measure of electric signal strength, magnetic recordings additionally provide a measure of magnetic field direction.

**Figure 1.**
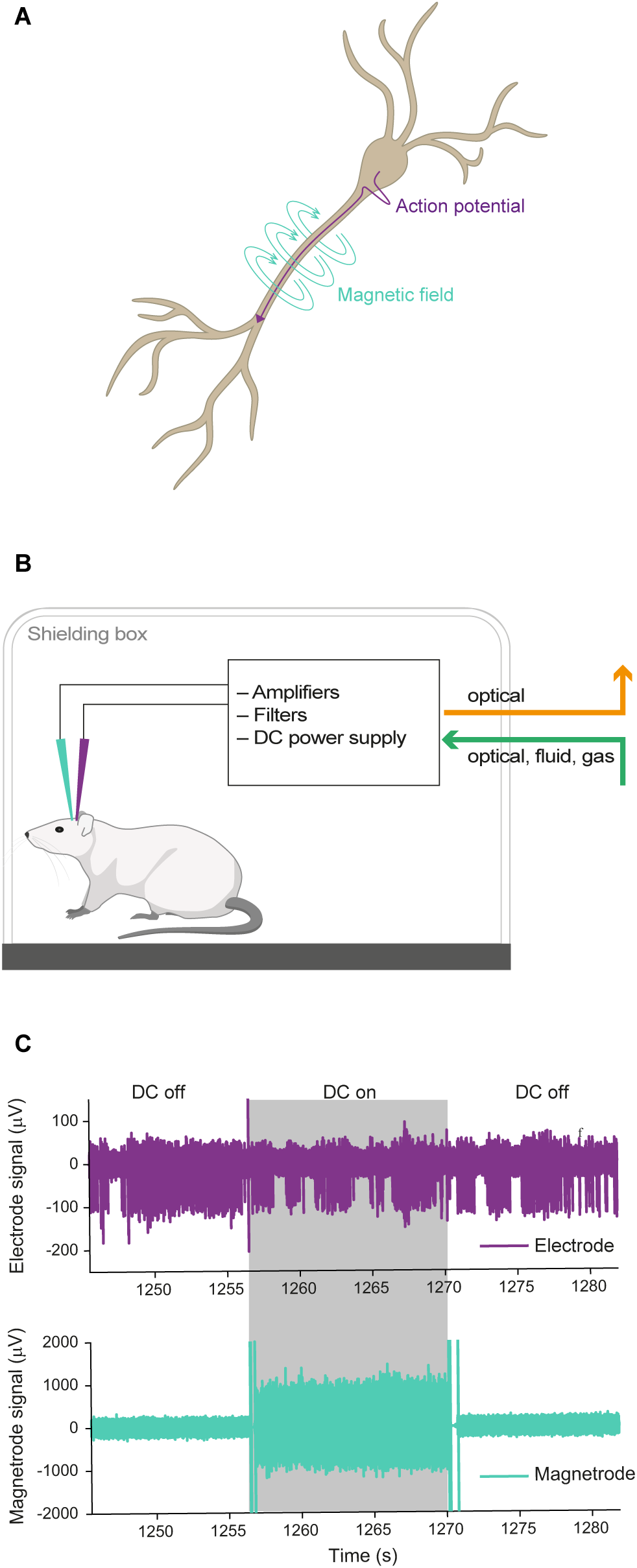
Biophysical background and recording setup. **A** Illustration of a pyramidal neuron and the electric currents (purple) and magnetic fluxes (turquoise) generated by an action potential originating at the soma and travelling along the axon. **B** Illustration of the recording setup. The anesthetized animal was placed in a Faraday cage. We recorded simultaneously with electric (purple) and magnetic (turquoise) sensors. All electronics necessary for the magnetic recording were contained within the Faraday cage, and no electric connections were entering or leaving the cage. **C** Example raw traces for the electric and magnetic sensors. Switches of the DC supply could occur pseudo-randomly every 10 seconds and are indicated here by shading (grey: DC on, white: DC off). A switch of the DC supply caused an artifact on both sensors. As expected, the DC condition did not affect the recorded signal on the electrode, while the GMR sensor can only detect magnetic signals in the DC on condition.

The fact that magnetic signals are mostly unaffected by tissue, combined with the fact that their recordings provide directional information, could be utilized to improve action potentials recordings. Action potentials generate currents primarily at the neuronal cell bodies, and dozens of such cell bodies surround any given sensor position in neuropil. Nevertheless, at a given position, a microelectrode contact typically provides spike waveforms of merely 1-10 single neurons (Buzsáki, 2004). Other neurons in the close vicinity of the recording electrode might not be detectable because they are too strongly electrically insulated by the dense mesh of cell membranes. These membranes are essentially transparent to magnetic fields, which might enable the simultaneous recording of large numbers of single neurons. The detectable neurons could then also be source localized by using the directionality of the magnetic signal. This would be aided by future probes that combine magnetic sensors with sensitivity in different directions in close spatial proximity.

Given these clear theoretical advantages, our goal was to record the magnetic signals of neuronal action potentials (APs) *in vivo*. Recording magnetic AP signals *in vivo* is technically very challenging. It is generally difficult to shield magnetic interference from external artifact sources. Furthermore, magnetic recordings require advanced technology. Conceivable approaches include (1) coils, which would however be difficult to miniaturize for AP recordings; (2) optically pumped magnetometers, which would suffer from the same problem; (3) nitrogen-vacancy centers in diamonds, which can be miniaturized sufficiently but require the application of both microwaves and polarized light; (4) giant magnetoresistive (GMR) sensors, which can be miniaturized and do not pose these challenges. Therefore, we used GMR sensors to investigate the feasibility of *in vivo* magnetic AP recordings. Achieving this requires the use of miniaturized magnetic sensors that can be positioned in the immediate vicinity of the neuronal cell bodies. GMR sensors provide sufficiently high sensitivity at a surface size of 30 x 30 μm. GMR sensors show a resistance that is proportional to the strength of the magnetic field and can yield a measurement sensitivity in the nanotesla (nT) range (Chopin et al., 2020). To measure the magnetic field, a small current is passed through the GMR sensor, and magnetic-field related resistance is measured as voltage across the sensor.

We have recently been able to record the magnetic counterparts of neuronal LFPs from the visual cortex of anesthetized cats using GMR sensors on silicon backbones, which we refer to as “Magnetrodes” (Caruso et al., 2017). However, these probes were too large to bring the magnetic sensors sufficiently close to intact, spiking neurons. Therefore, we further miniaturized the probes to be located close to the tip of 25-micron thick silicon needles, similar to typical silicon-based multi-contact electrodes (Chopin et al., 2020). Here, we used conventional electrophysiological techniques to record electrical neuronal spikes in anesthetized rat hippocampus, in combination with two high-sensitivity, miniaturized GMR sensors to uncover the magnetic counterparts of single-neuron action potentials in vivo. This is a proof-of-principle study to evaluate the possibility of recording magnetic signatures at the level of single unit action potentials.

## Methods

### EXPERIMENTAL MODEL AND SUBJECT DETAILS

#### Animals

A total of 8 male Sprague-Dawley rats (Janvier Labs, France) were used in this study (4 for the first set of experiments, 4 for the second set). Rats were all approximately 7 to 8 weeks of age (330-420 grams). We used only male rats as they are bigger at this age, which was advantageous for the experiments. Animals were housed in pairs or groups of 4 animals. All animal experiments were in accordance with the German law for the protection of animals and the “European Union’s Directive 2010/63/EU” and approved by the local government office (Regierungspräsidum Darmstadt).

### METHOD DETAILS

#### Surgical procedures

All experiments were performed under general anesthesia. The rat was anesthetized with an injection of Ketamine (80 mg/kg) in combination with Medetomidine (0.01 mg/kg). Anesthesia was maintained throughout the experiment with Isoflurane (0.5% – 2% in 100% oxygen). For analgesia, the animal received Buprenorphine (0.3 mg/kg) subcutaneously (s.c.) every three to five hours. The animal was placed in a stereotaxic frame (Kopf Instruments, USA) and received a s.c. injection of Dexamethasone (1 mg/kg) to prevent edema. Every two to three hours they also received a 2 ml bolus injection of a solution containing amino acids and glucose (Aminomix 1 Novum, Fresenius Kabi). Heart rate, respiration rate and body temperature were continuously monitored throughout the experiment.

The skin was removed over the skull, and a craniotomy was performed centered on 4.5 to 5 mm posterior of Bregma and 4 mm lateral to the midline. To facilitate probe insertion, we opened a slit in the dura mater using a manually bent hypodermic needlea (Sterican, 30G). The probes were lowered either into the cortex or to a depth of approximately 2 mm to record in the hippocampus. At the end of the experiment, the animal was euthanized using an overdose of Narcoren (min. 160 mg/kg).

#### Probes

##### Magnetic Sensors

The magnetic sensors or “Magnetrodes” used in this study are devices for local magnetic sensing based on spin electronics principle, where the electrical transport varies as a function of the magnetic configuration of a set of very thin (nanometers) magnetic layers.

The Magnetrodes used here have been produced by depositing the magnetic layers on a SOI substrate, allowing a thinning process where the tip of the probe, which contains 2 magnetic sensing elements, is reduced down to 25μm to facilitate insertion within tissues. Details on the probe fabrication can be found in Chopin et al. (2020). The Magnetrode’s sensing elements use the Giant Magneto Resistance principle (Baibich et al., 1988; Dieny et al., 1991) effect. They are designed as a meandering structure of 5 segments, each of which is 4μm wide and 30μm long. The two GMRs are separated center to center by 250μm. The magnetic sensors are kept electrically decoupled from the environment by a Al2O3 (150 nm)/Si2N4 (150 nm) passivation bilayer.

Prior to the experiments, magnetotransport and noise measurements were performed to characterize the GMR sensors. Their sensitivity is in the range of 1.5-2%/mT and their limit of detection (i.e. the signal amplitude at which the signal-to-noise ratio is equal to 1) at 1 kHz of 1nT/√*Hz*. To allow for continuous recording including experimental and control condition, we set up a system in which we switched the bias voltage to the GMR sensor on and off in a pseudorandom way, thereby interleaving DC on and DC off blocks. Every 10 seconds during a recording, a pseudorandom decision was made to either change the DC state or to leave it as it was before. This led to at least 10 s long blocks per condition. Over a continuous recording block, the number of 10 s DC on and DC off blocks was equal.

##### Electrodes

In the first round of experiments, we used Tungsten microelectrodes (FHC, USA) to record the electrical spikes as ground truth. One or two Tungsten electrodes were manually attached to the Magnetrode under a microscope. The tip of the microelectrode was placed close to the lowest GMR sensor. In the case when two microelectrodes were used simultaneously, they were positioned with a vertical offset. In the second set of experiments, the Tungsten electrodes were replaced by multi-contact silicon arrays (A1×32-Poly2_HZ32 & A1×32-Poly3_HZ32, NeuroNexus, USA). These have the advantage of allowing for wider coverage and denser sampling of the surrounding tissue, making it possible to spike-sort the data. They were glued to the Magnetrode in a similar way as the Tungsten electrodes, with their contacts facing outward.

#### Acquisition of electric and magnetic signal

The animal was placed inside a Faraday cage (Fig 1B). The Faraday cage was built in house and consisted of a single layer of aluminum, providing weak magnetic shielding. This explains the relatively high noise floor in the magnetic recordings and could be improved in future recordings. For reference, Figure 2 shows power spectra for the magnetic sensors in the recording situation in the brain (Fig 2) and measured in air in a well-shielded room (Fig 2, note that the peak at 30Hz is due to a local calibration coil emitting a signal at that frequency). All connections going in or out of the Faraday cage used here were optical, apart from a gas and a water line for the anesthesia and the water bed used to keep the animal warm. This was done in order to reduce electric and magnetic noise for the recordings. Electric signals were recorded via active, unity gain headstages (ZC32, Tucker Davis Technologies, USA) and digitized at 24,414.0625 Hz or 48,828.125 Hz (PZ2 preamplifier, Tucker Davis Technologies, USA). The TTL signal controlling the DC on/off switches was recorded at 48,828.125 Hz. Each GMR sensor was measured in a Wheatstone bridge configuration with a variable resistance for adjustment (Chopin et al., 2020), fed on DC bias. Low noise amplification of the output voltage was performed with a Texas Instrument INA 103. A second stage of filtering amplification (0.3-30 kHz) of the GMR signal was obtained with a low noise amplifier (SR 560-Stanford Research Systems). To reduce noise and avoid contamination by 50 Hz power line signals, the bias voltage and both levels of amplification for the GMRs were battery powered.

**Figure 2.**
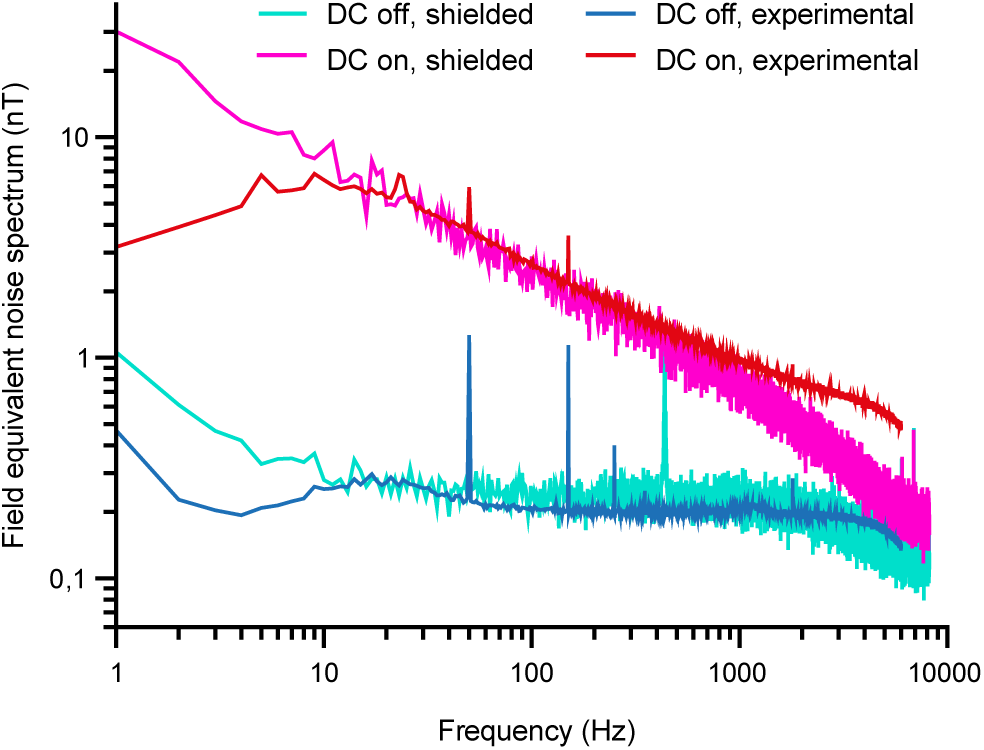
Power spectra of the GMR sensor. Example power spectrum from one in vivo (minimally shielded) recording block and for a control recording with a similar sensor in air in a well shielded room. The experimental condition spectrum shows peaks for electric line noise at 50 Hz and 150 Hz, and additional noise peaks at 5, 9, and 23 Hz of unknown origin in the DC on condition. The peak at 30 Hz in the well shielded room is due to a local coil used for calibration emitting a signal at that frequency. Note that the axes are in logarithmic scales.

### QUANTIFICATION AND STATISTICAL ANALYSIS

All analyses were performed using custom MATLAB (MathWorks) code.

#### Spike detection

For the thresholding approach, we first band-pass filtered the electric signal using a fourth-order Butterworth filter (500 – 5,000 Hz). In the filtered signal, peaks were detected (*findpeaks* function of MATLAB) for each electrode. Peaks that crossed a pre-set threshold were identified as spikes and used for further analysis. For the spike sorting approach, we used the “Kilosort” algorithm (Pachitariu et al., 2016) or, in the case of the Tungsten data, the “spyKING CIRCUS” toolbox (Yger et al., 2018). For the single units recorded on Tungsten units, a manual inspection after the algorithm step was performed. Well-isolated single units were identified by inspection of the wave shape, the ISI distribution and the number of events detected. Since we only had two recording channels available in these recordings, some of the clusters identified by the algorithm did not resemble a neuronal wave shape and were most likely clusters of noise artifacts and thus not suitable for further analysis.

#### Processing of magnetic signal

The magnetic signal was filtered using a fourth-order Butterworth filter (500 – 5,000 Hz) for the analysis based on thresholded spikes, and using a second-order Butterworth filter (5 – 6,000 Hz) for the analysis based on spike sorting. The DC switch between DC-on and DC-off blocks induced artifacts in all signals (illustrated in Fig 1C). Therefore, we discarded the first 2 seconds after a switching event as well as the last 0.5 s before a switch from both the electric and the magnetic signal.

#### Spike triggered average

To calculate the spike triggered averages, we first determined for every detected spike whether it occurred during a DC on or a DC off block. All spikes that fell into the artifact period around a switching event were ignored. For each condition individually, we centered a window on each detected spike and cut these windows from the continuous signal. We then averaged these windows per condition using a trimmed mean to reduce the influence of outliers (*trimmean10* function of Matlab). For the spike-sorted data, we performed this analysis per cluster, for the thresholded data for all spikes detected on a given electrode.

For the electric STAs based on spike sorted units, the raw electric signal was filtered using a second-order Butterworth filter (5 – 6,000 Hz).

#### Correlation analysis

We extracted the central 2 ms of each electric STA and its corresponding magnetic STAs. In a next step, every STA was corrected for its mean. We then calculated individual cross correlation values for the electric STA with the magnetic STA per sensor. For the cross correlation, we shifted the electric signal sample-by-sample across the magnetic signal. Each shift corresponded to 1 sample (sampling rate 24,414.0625 Hz, 1 sample ≈ 0.04 ms), and the maximal shift was +/-10 samples. Significance was assessed for each correlation value individually and Bonferroni-corrected for the number of sample shifts, single units, and sensors.

#### Estimation of magnetic signal strength

To estimate the magnetic peak-to-peak amplitude, we subtracted the magnetic STA of the off condition from the magnetic STA of the on condition, such that the capacitive coupling present in both conditions is subtracted out and the remaining signal can be assumed to be purely magnetic.

To test whether the measured difference was bigger than what could be expected by chance, we performed a min/max based permutation test (Westfall and Young, 1993). We ran 1,000 difference calculations in which the trials were randomly assigned to the on- or off-condition, keeping the number of trials per condition consistent with the observed values. From each resulting difference, we saved the minimum and maximum values across all time points. The observed difference was then tested against the 97.5^th^ percentile of the maximum distribution and against the 2.5^th^ percentile of the minimum distribution. This corresponds to a two-sided test at an alpha level of 0.05, corrected for the multiple comparisons across time points.

To transform the measured μV values into nT we use the sensitivity of the probes as experimentally determined under a known magnetic signal (see Chopin et al. (2020)), defined as the voltage variation at the bias voltage of the GMR (here 1 V) per field unit and is expressed in V/T.

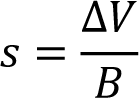

The output voltage needs to be divided by the amplification gain in the acquisition chain (here 1000), so the output signal in T is:

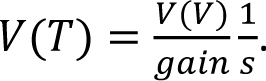

With a sensitivity of 18.5 V/T, 100 μV signal corresponds to 5.4 nT.

## Results

### Spike triggered averages based on thresholded data reveal noise artifact

We simultaneously recorded electric and magnetic signals from the brain of anesthetized rats placed in a Faraday cage (Fig 1B,C). By switching the DC supply to the GMR sensors on and off at pseudorandom times during the experiment, the recordings included randomly interleaved control blocks during which the GMR sensor did not receive any external current. Periods containing the artifact that was introduced by the switching of the DC supply were removed during data processing. In a first set of experiments, we used Magnetrodes with Tungsten electrodes positioned in close proximity to one of the magnetic sensors (Fig 3A). With this probe combination, we recorded data either from visual cortex or from the hippocampus underneath the visual cortex. High-amplitude spike events were extracted from the continuously recorded data by detecting peaks and selecting those that exceeded a pre-determined threshold. The spike-triggered average (STA) of the magnetic signal around these events revealed a signature in a subset of recordings. Note that all magnetic STAs were kept on the μV scale of the electric recording system, because it is possible that the data contains a mix of magnetic and electric signals, due to capacitive coupling (see below). Fig 3B shows STA results for one example recording. In this example a threshold of 5 SDs was used to detect spike events. At this threshold, we detected 15.119 events. A window centered on the time of each detected spike events was cut from the continuous electric and magnetic recordings. These windows were averaged per condition. The resulting STA for the DC-on condition is illustrated in red, for the DC-off condition in blue. The electric STA indicates, as expected, no difference in the DC-on and the DC-off conditions. For the two magnetic sensors, a peak appeared in the DC-on condition, corresponding to the condition in which the GMR sensor is sensitive. However, when we lowered the threshold for spike detection, this signal became bigger (left two columns Fig 3C), and when we increased the threshold, the signal decreased and even disappeared in some instances (right two columns Fig 3C). The electric STA was also affected by changes in the threshold, however becoming more pronounced with higher thresholds (top row Fig 3C – note the changing y-axis scales for Fig 3C).

**Figure 3.**
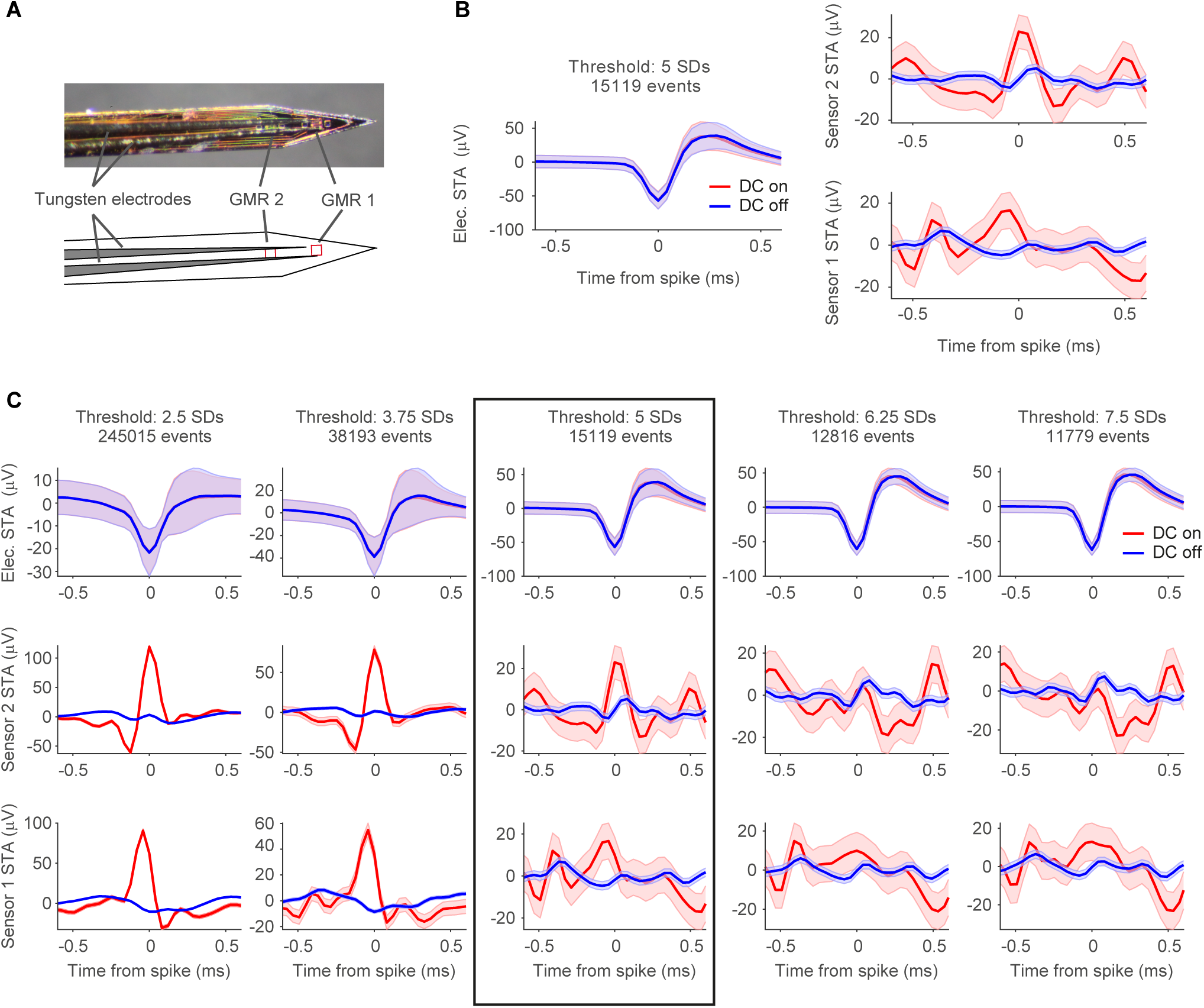
STAs on thresholded spikes. **A** Photo of the magnetic probe with two GMR sensors and one Tungsten electrode glued onto it, and schematic drawing of the probe arrangement below the photo. **B** Electric and magnetic STAs of one example recording session, with spikes detected at a threshold of 5 SDs (15.119 spike events detected at this threshold). Left plot shows the STAs resulting from triggering the continuously recorded electric signal on the thresholded spikes from the same electrode. Right plot shows the STAs of the signals recorded from the two magnetic sensors, respectively. All STAs were calculated separately for the two recording conditions (red: DC on, blue: DC off). The STA on magnetic sensor 2 (top right) shows a peak around the time of the spike only for the DC-on condition. Shaded area represents standard error of the mean. **C** Same recording as in B, now with varying thresholds. From left to right the threshold is increased from 2.5 SDs to 7.5 SDs in steps of 1.25 SDs. The top row shows the electric STAs, the middle row the STAs for magnetic sensor 2, bottom row for magnetic sensor 1. The threshold setting illustrated in B is marked with a black box. From left to right, a decrease in the peak amplitude can be observed for the two magnetic sensors, while the peak in the electric STA is becoming more pronounced. Shaded area represents standard error of the mean. Please note that the y-axis scales are different for the different threshold settings.

This tight relationship to the selected threshold for both, the electric and the magnetic STA, suggests that the signature in the magnetic STA arises due to an external artifact rather than a magnetic signal. Such an artifact could be induced by correlated electric and magnetic noise from an external source being picked up by both, the magnetic and the electric sensors.

### STAs based on isolated single units reveal magnetic signature

One way to avoid this artifact is to separate spikes from noise by applying state-of-the-art spike-sorting algorithms. These sorting algorithms utilize template matching of the spike waveforms rather than relying purely on amplitude thresholds, thus reliably separating spikes of putative single neurons, commonly referred to as single units, from noise events (Pachitariu et al., 2016; Yger et al., 2018).

We first selected the recording block with the most robust peak in the magnetic STA of the thresholding analysis (Fig 4A) and identified two well-isolated single units after spike-sorting (Fig 4B, D). Both units showed a pronounced deflection in the magnetic STA on magnetic sensor 1 (Fig 4C, E). The amplitude of this signal was much larger in the DC-on condition in which the GMR sensors are active. Note that there are differences in the number of spikes when comparing the DC-on and DC-off conditions, which might be due to a small increase in temperature of the surrounding tissue caused by the electric current in the GMR sensor. Since the spike counts for each unit are the results of a spike-sorting procedure, the difference cannot be attributed to artifacts.

**Figure 4.**
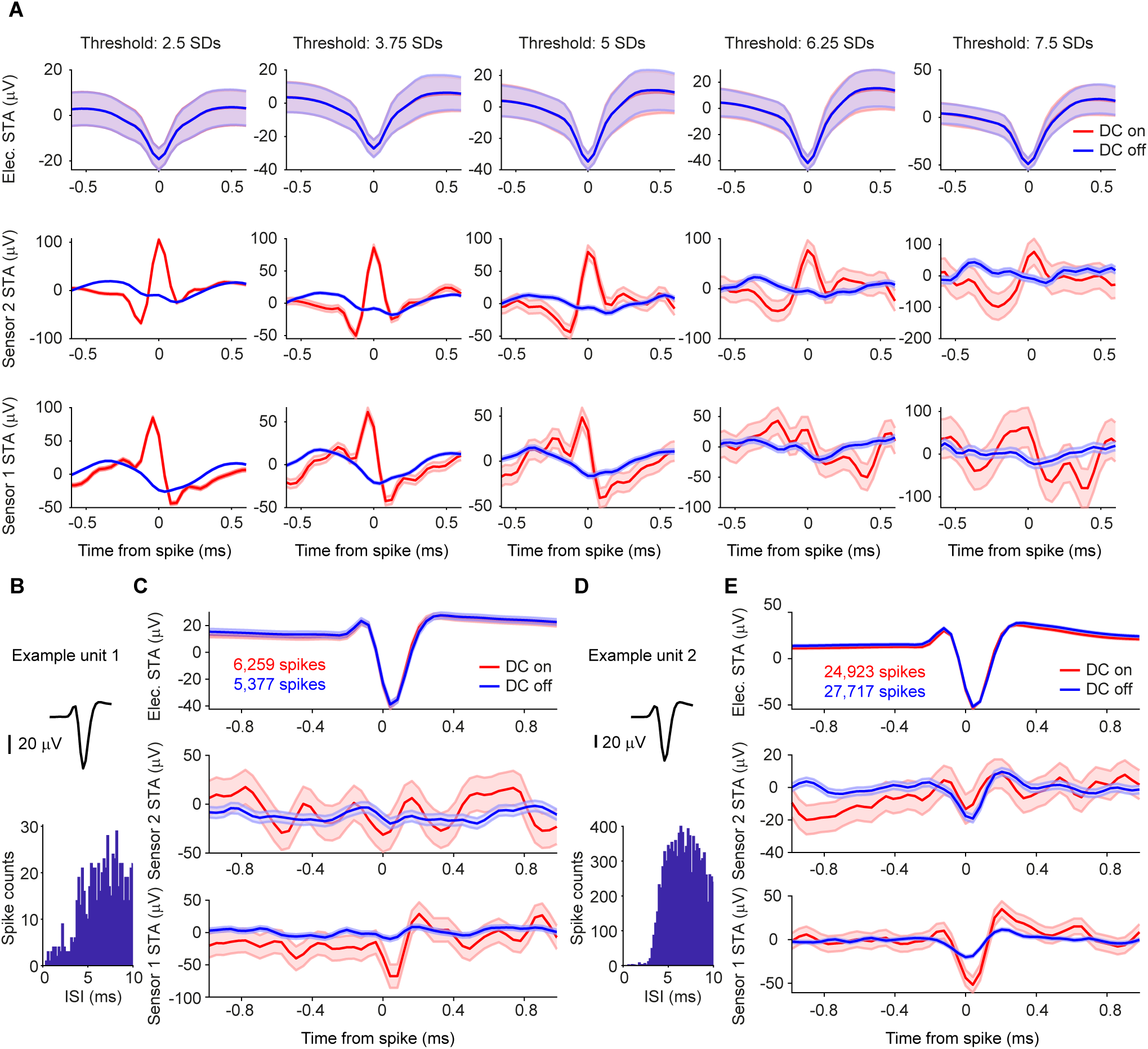
STAs on thresholded versus spike-sorted data from Tungsten probes. **A** Electric and magnetic STAs for one recording block. As the threshold increases from left to right in steps of 1.25 SDs, like for the example shown in Fig 2, the electric STA (shown in the top row) becomes more pronounced, and the magnetic STA on sensor one (in the bottom row) disappears. The peak on sensor 2 (middle row) is less affected and still visible at higher thresholds. Shaded area represents standard error of the mean. Please note that the y-axis scales are different for the different threshold settings. **B** Average waveform and inter-spike interval (ISI) distribution for one example unit after spike-sorting the block shown in A. **C** Top row: STA of the continuous data recorded on one of the 2 Tungsten electrodes triggered on the spikes of example unit 1 per condition (red: DC on, blue: DC off). The number of spikes detected per condition is given in the inset. Shaded area represents standard error of the mean. Middle row: STA of the magnetic signal recorded on the magnetic sensor 2 per condition. The higher level of background noise in the DC-on condition is expected. Bottom row: same as middle row but for magnetic sensor 1. A clear magnetic signature can be observed on sensor 1 only in the DC-on condition. **D, E** Same as B, C, but for example unit 2.

A total of 12 blocks were recorded using a Magnetrode with 2 Tungsten electrodes attached to it. We spike-sorted all 12 blocks and identified a total of 31 well-isolated single units (see Methods). To assess whether a single unit exhibited a magnetic spike signature on one of the adjacent GMR sensors, we calculated the cross-correlation between the magnetic and the electric STAs for the central 2 ms around the time of the electric spike. We investigated sample-by-sample shifts for up to 10 samples in each direction (1 sample ≈ 0.04 ms), since the time course of the magnetic spike could differ from that of the electric spike (Fig 5A for illustration). For each cross-correlation, we identified the peak correlation value and tested this correlation for significance (using Bonferroni correction for multiple comparisons across units, magnetic sensors and sample shifts). Using this approach, we identified 4 single units with significantly correlated signals on the two sensor types (Fig 5B). Two units showed this correlation for the magnetic signal recorded on sensor 1, one unit for the signal recorded on sensor 2, and one unit showed a correlation with the signal on both magnetic sensors. Note that the correlations tend to show a peak (maximal y-value) for slightly positive shifts (x-values), which suggests that the magnetic signal is leading the electric spike. A detailed consideration of this shift is provided in the discussion.

**Figure 5.**
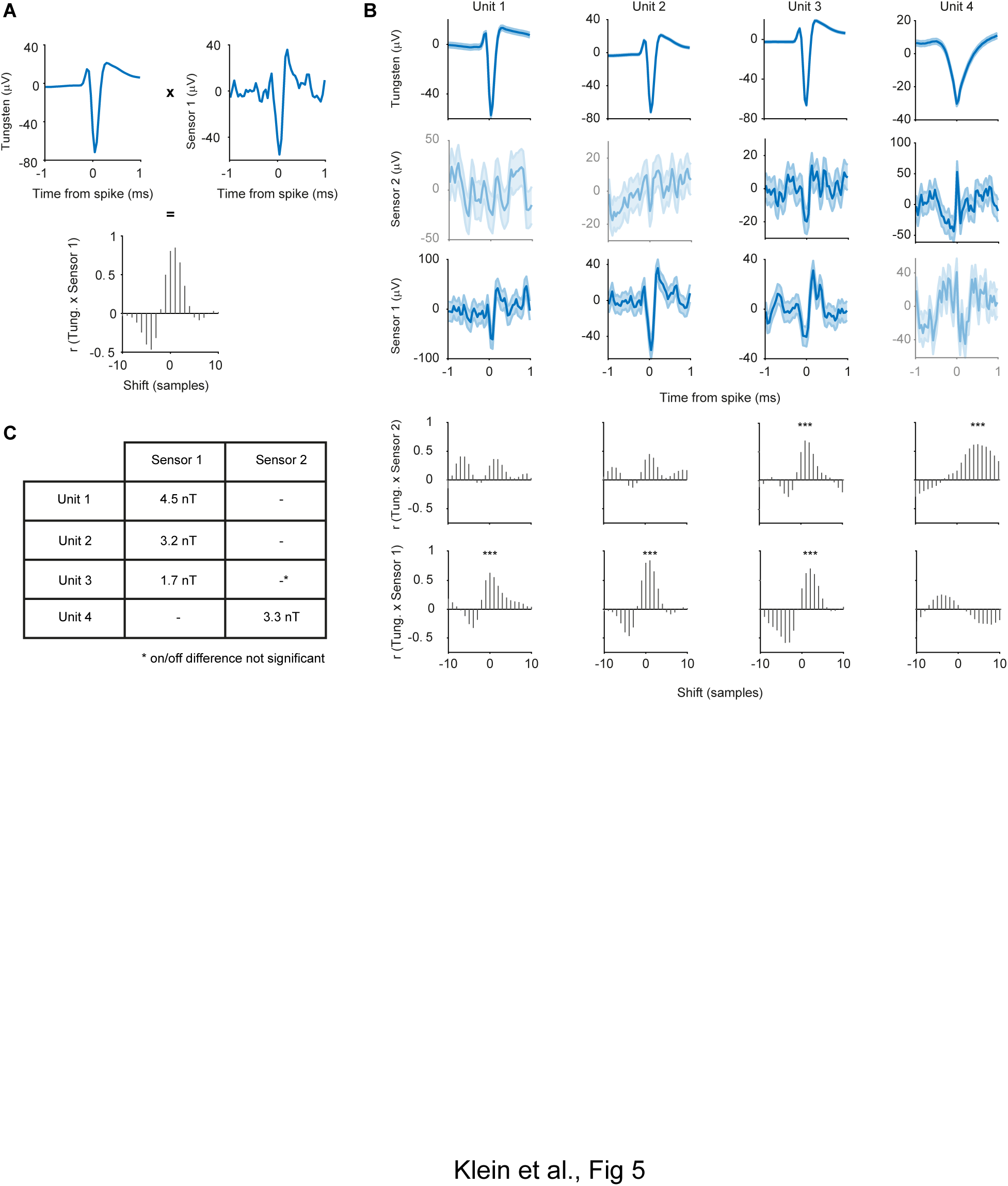
Correlation analysis of electric and magnetic waveforms. **A** Illustration of the correlation analysis. Top left plot shows the demeaned central 2 ms of the electric STA of one well isolated single unit. The top right plot shows the corresponding demeaned magnetic STA on sensor 1. We computed the cross correlation values for shifts of +/- 10 samples around the time of the spike of the electric STA with respect to the magnetic STA. The resulting distribution of correlation values can be seen in the bottom plot. **B** Tungsten and magnetic-sensor signal, as well as cross-correlation values for the four significantly correlated single units. Shaded area represents standard error of the mean. Not significantly correlated signals are shown at decreased intensity. Unit 2 is the example used in A. Significant correlations are marked by asterisks. **C** Table of signal strength for the significant magnetic signals. The significant correlation with sensor 2 for unit 3 does not differ from chance in amplitude when tested for on/off condition difference.

The STA analysis also revealed a small deflection in the DC-off condition (Fig 4E). This most likely indicates capacitive coupling. Capacitive coupling refers to currents in the tissue inducing currents in the electric circuits of the GMR sensor. The electric contacts of the GMR constitute a capacitor that is isolated by an insulation layer from the surrounding bath. If the insulation were perfect and the voltage in the bath were homogeneous, the electric signal would not be visible on the GMR. However, since the neuronal signal source likely generates an inhomogeneous voltage field, a capacitive coupling signal can appear on one or both sensors. The resulting artifacts reflect the ongoing current fluctuations in the tissue and can thus resemble a spike-like fluctuation in the GMR signal. Since capacitive coupling decreases with lower frequencies, it might be of less concern and influence when recording LFP signals (as in our previous study Caruso et al. (2017)). At higher frequencies, necessary for AP detection, the capacitive coupling is expected to be more apparent. In order to be able to estimate the amplitude of the true magnetic signal, we subtracted the STA in the DC-off condition from the STA in the DC-on condition. Capacitive coupling should be present in both conditions, independent of the bias voltage, hence not amplified in the DC-on condition, and therefore will be eliminated by this subtraction, because the sensor is only able to measure a true magnetic signal when the DC supply is turned on. The experimental design with multiple DC-on and DC-off blocks randomly interleaved in each recording session provided optimal conditions for this subtraction, because the sensor location and the sources of the recorded spikes did not change between the two conditions. The differences on the significantly correlated channels were bigger in amplitude than would be expected by chance (see Methods) for all 4 units. However, unit 3 (Fig 5B), which was significantly correlated with the magnetic signals on both sensors, showed a significant on-off difference only on sensor 1. Since we can now assume the signal to be purely magnetic in nature, we transformed the μV scale into a nT scale (Fig 5C).

Tungsten electrodes often yield only a very small number of sortable single units. To increase the yield of single units with a high number of detected events, we switched to 32-channel silicon probes mounted on the Magnetrode (Fig 6A) for our second set of experiments.

**Figure 6.**
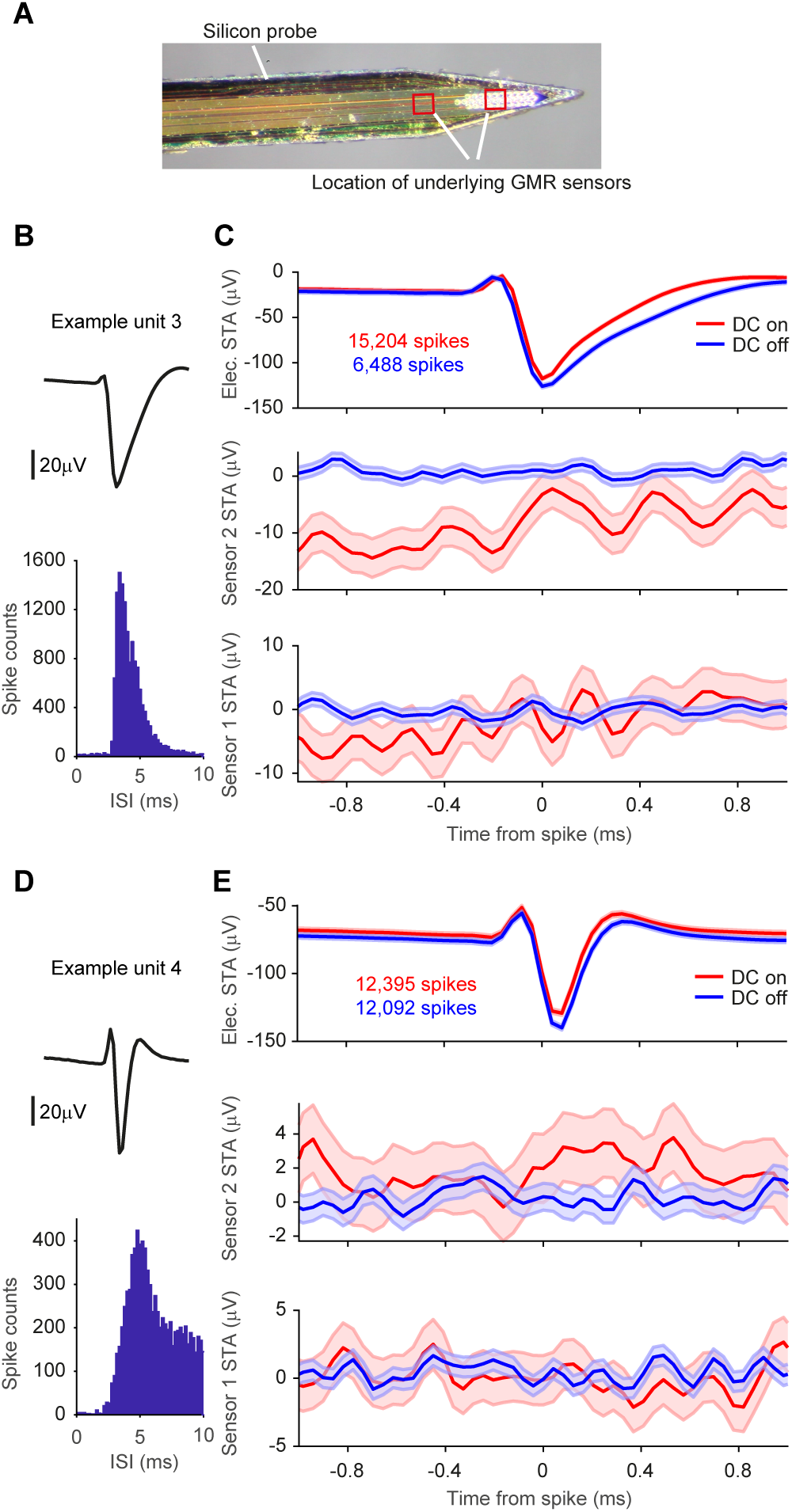
STAs on spike-sorted data from silicon probes. **A** Photo of the magnetic probe and the silicon probe with 32 electric contacts glued onto it. Locations of the underlying GMR sensors are indicated with red boxes. **B** Average waveform and inter-spike interval distribution for one example unit. **C** Top row: STA of the continuous data recorded on one of the 32 electric channels triggered on the spikes of example unit 3 per condition (red: DC on, blue: DC off). The number of spikes detected per condition is given in the inset. Shaded area represents standard error of the mean. Middle row: STA of the magnetic signal recorded on the magnetic sensor 2 per condition. The higher level of background noise in the DC-on condition is expected. No apparent magnetic signature can be observed for either condition. Bottom row: same as middle row but for magnetic sensor 1. **D, E** Same as B, C, but for example unit 4.

Recordings using this probe combination were performed in hippocampus and yielded a large number of single units after spike sorting (n = 593 from 7 recording blocks). Fig 6B and D show two example units. Calculating magnetic STAs based on individual single units did not show any apparent magnetic signature (Fig 6C and E).

We focused our further analysis on well-isolated single units with a high number of events. We considered a single unit to be well isolated if less than 1% of spikes occurred within 2 ms of the preceding spike (reflecting the refractory period of the neuron) and excluded units with less than 7000 spikes. Across the 7 recording blocks, we found a total of 73 of such well-isolated single units (73 units out of 593 units, corresponding to 12%, minimum number of spikes in the distribution = 7,038, maximum number of spikes = 279,872, 25^th^ percentile = 10,974 spikes, 75^th^ percentile = 31,328 spikes). None of the 73 well-isolated single units showed a significant correlation with the magnetic signal on either sensor (Bonferroni correction for multiple comparisons across units, sensors and sample shifts). Thus, while the switch from Tungsten electrodes to silicone probes strongly increased the number of recorded single units, no magnetic signatures were detected in this configuration.

### Increasing the sensitivity of the GMR sensor leads to heating artifact

In an attempt to increase the yield of magnetic AP signatures, we increased GMR sensitivity by increasing the voltage applied across the sensor. We recorded two blocks in which 2 V instead of 1 V were applied to the GMR. However, these data revealed that at this voltage the heating artifact mentioned earlier (see consideration about differences in spike count above) substantially affected the spiking behavior of the recorded neurons, not only in spike rate but also in spike shape. The effect of temperature on neuronal activity has been reported before (see for example Hedrick and Waters (2011), Hedrick and Waters (2012)). In our case, it leads to a disturbance in the waveforms of the electric spikes and increased the spike count in the on condition compared to the off condition. The extent of heat-related disturbance in the 2 V blocks made direct comparison of the two conditions problematic and hindered the reliability of spike sorting results. We therefore decided to exclude those data from the analysis.

## Discussion

After having previously demonstrated that it is possible to record the magnetic signatures correlated to ERPs *in vivo* (Caruso et al., 2017), we performed two sets of experiments in an attempt to measure magnetic signatures of APs *in vivo*. Our initial approach of using simple amplitude thresholds to detect electric spikes revealed the importance of reliable spike sorting methods in order to avoid artifacts caused by correlated electro-magnetic noise. With the refined spike sorting approach, we identified four single units with significant magnetic signatures around the time of the electric spike. The waveform of this magnetic signature was highly correlated with the waveform of the electric signal and was significantly bigger in the DC-on than the DC-off control condition for all four units. Since the number of single units that can be identified using two Tungsten electrodes is low, we increased the single unit yield by performing additional experiments with silicon electrode arrays mounted directly onto the GMR sensors. We were able to record a large number of single units with this arrangement. However, none of the units recorded with silicon probes showed a magnetic spike signature that was statistically significant. When increasing the sensitivity of the GMR by increasing the voltage applied to it, we observed a clear heating artifact and had to exclude the data recorded with this setup from further analysis.

We obtained magnetic signatures of electrically recorded spikes only from a small proportion of the units recorded with Tungsten electrodes and from none of the units recorded with silicon probes. The silicon probes were attached to the Magnetrode and thereby created an obligatory distance to a neuron of at least the probe thickness (15 μm). This distance was orthogonal to the main orientation of neuronal dendrites in the recorded areas. The distance might have prevented successful magnetic recordings. Using silicon probes in different spatial configurations with respect to the magnetic sensor could potentially avoid this problem and is an interesting option for future research. By contrast, the Tungsten electrodes had conical tips that left a gap between the Tungsten tip and the magnetic sensors. In this gap, neurons could come in very close proximity to the magnetic sensors. It is even conceivable that neurons were “trapped” in this gap during probe insertion. Note that in one case, a magnetic spike signature was detected on sensor 2, which was 250 μm above sensor 1 with the immediately neighboring Tungsten tip, but this separation was parallel to the main orientation of neuronal dendrites in the recorded areas. The pattern of results is consistent with successful magnetic recordings requiring that the distance between the neuron and the magnetic sensor be very small. Even at this distance, averaging of many action potentials, based on simultaneous electric recordings, was required to reveal the magnetic signature. Further improvements in sensitivity will be required to enable the direct magnetic recording of individual action potentials in vivo. Yet, our approach demonstrates a proof of principle and it provides a measurement of the amplitude of the magnetic action potential, which is crucial for further developments in this area.

We observed that magnetic spikes tended to show a very slight temporal lead over electric spikes, and that the capacitive component during DC-off tended to show a very slight temporal lead over the magnetic component during DC-on. Overall, the observed shifts were very small, in the range of one sample, corresponding to 40 microseconds. There are a number of factors that could generate or influence these shifts: (1) The electrode has an impedance spectrum that generates phase delays and thereby time delays. (2) The magnetrode signal is recorded via an electronic circuit that generates small, but non-negligible time delays. (3) The magnetic signal on the one hand and the electric and capacitive signal on the other hand reflect different underlying processes. The magnetic signal most likely reflects primary intracellular ionic currents, occurring after the AP has emerged at the AP-initiation zone and invades the cell body. The electric and capacitive signals most likely reflect extracellular return currents. The return currents are thought to occur simultaneous to the primary intracellular currents, yet they are expected to have a different spatio-temporal profile. Furthermore, the magnetic versus the electric recordings (and to a lesser extend the magnetic versus the capacitive recordings) are expected to have slightly different spatio-temporal integration profiles. These factors together can lead to small time shifts between magnetic versus electric or capacitive signals. These considerations are also relevant for the question of whether the observed magnetic signal has a dipolar or quadrupolar source. We consider it most likely that we measured a dipolar source. Dipolar sources arise in the cell body, and thus from the biggest magnetic field. Quadrupolar sources are mainly limited to the axon and/or the axon initial segment with the AP-initiation zone and are probably too small to be recorded by our sensors. Additional support for this argument is provided by the fact, that the axon and the axon initial segment are approximately rotationally geometric, which would further cancel quadrupolar field contributions.

A previous attempt at measuring APs using magnetoresistance in slices from mouse brain provided promising preliminary results (Amaral et al., 2011). However, in these experiments, it was not possible to separate a putative magnetic signal from potential capacitive coupling. To achieve this separation, several subsequent studies using magnetoresistance, including the present one, used different approaches. Barbieri et al. (2016) APs in a muscle-nerve preparation and used a control for capacitive coupling that was possible in this specific setting: they turned the muscle fiber by 90 degree, which is expected to rotate the magnetic signal out of the direction of sensitivity of the magnetic probe. Caruso et al. (2017) used a high-frequency (20-80 kHz) modulation-demodulation approach to successfully separate a capacitive component from the visually evoked field. The signals measured in these preparations were within the range predicted and estimated for *in vivo* recordings in the central nervous system (Petridou et al., 2006; Hall et al., 2012). Finally, here, we used a new approach enabled by new technical developments that allowed us to record interleaved DC-on and DC-off conditions. The DC-off condition provides a clear estimate of the capacitive component, which was much lower than observed in the study of Amaral et al. (2011). While our approach was able to identify a magnetic AP signal that exceeded a putative capacitive component, and while it was able to measure AP signals from isolated single neurons, it was not (yet) able to measure the magnetic counterpart of single APs. Rather, we needed to use the simultaneously recorded electric spike signals to trigger the averaging of the magnetic signal and thereby reveal the magnetic AP recordings. For the empirical data, we used a threshold of at least 7000 electric spikes for this averaging process. We also calculated the theoretical limit for this number, given the observed noise spectra (Fig 2). In the frequency range up to 1 kHz, the noise spectra show a typical 1/f pattern, and the probe’s intrinsic noise is around 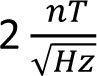, which corresponds to a detection level of 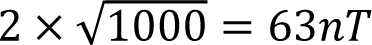. In the frequency range between 1 and 6 kHz, the noise spectra are at the thermal noise of 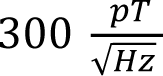, corresponding to 21 nT, such that the detection level is at 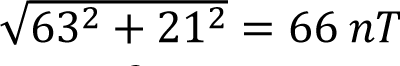. This means that to measure a signal of 1nT with an SNR of 1, one needs to have 66^2^ = 4356 acquisitions.

Magnetic recordings of action potentials from isolated axons have been achieved with nitrogen-vacancy (NV) centers in diamonds placed in close proximity to the squid giant axon or a large worm axon (Barry et al., 2016). However, this imaging technique is currently still limited to sensor placement outside of the examined tissue rather than within it. By contrast, our miniaturized GMR sensors can be introduced into nervous tissue, very similar to standard silicon-based electrodes. Interestingly, the peak-to-peak magnetic signal amplitude measured with NV diamond sensors from squid giant axons or isolated large worm axons placed directly onto the sensor were in the range of 4 nT. This is within the range of peak-to-peak amplitudes that we measured *in vivo*. Murakami and Okada (2006) predicted current dipoles in the order of 1 pAm; at 10 μm distance; for a 100 μm long neuron compartment, this would lead to around 200 pT, and for a 10 μm long structure, like the soma, 2 nT would be reached. Since the giant squid axon is larger in size than a neuron, it is theoretically expected to give a larger amplitude signal than the neuron. Yet, Murakami and Okada (2015) demonstrate that current dipole moment density is invariant across brain structures and species. Furthermore, by introducing the sensor into the tissue, we might have brought it closer to the source than previous studies (Barry et al. (2016) report 1.2 mm from the center of the axon in the intact worm, 300 μm in the excised worm). The greater proximity might have compensated for the smaller source size. A variety of parameters of the experimental paradigm and experimental conditions will influence the magnetic amplitude (Roth and Wikswo, 1985) and might explain some differences between theoretically predicted and empirically observed signals. Those parameters could not be measured in our *in vivo* recordings. For example, we do not know the precise spatial relation between the sensor and the signal source, and we do not know whether the spike signal originated fully or partly from the axon (initial segment) or the cell soma. These parameters might have been particularly favorable in the subset of our Tungsten-electrode recordings that led to detected magnetic spikes, and might thereby have contributed to relatively large observed magnetic field strengths.

As a non-invasive method to map magnetic fields induced by neuronal firing, Xiong et al. (2003) used a special acquisition technique in a conventional MRI scanner. They succeeded to greatly improve the temporal accuracy when scanning human subjects. However, a later study was not able to replicate this MRI approach to map neuronal activity (Parkes et al., 2007). Petridou et al. (2006) also used MRI scanning methods to measure activity in neuronal cell cultures. They were successful in detecting magnetic signals of neural activity. Additionally, new scanning paradigms have been suggested to use MRI to record magnetic signatures of ongoing neuronal activity *in vivo* (Sundaram et al., 2016; Truong et al., 2019; Roth, 2023). Yet, these measurements cannot be achieved at single neuron resolution, and the temporal constraints of MRI measurements make it nearly impossible to detect magnetic signature of single spikes.

A recent publication (Waterstraat et al., 2021) used MEG to record single-trial population spikes in humans. While this is an impressive extension of the MEG technology, it is not possible to perform this type of measurement at the level of spiking of individual neurons. Thus, with regard to magnetic recordings of action potentials from single neurons, the main candidate technologies are NV diamonds and GMR. For GMRs, this study provides a proof of principle for such recordings *in vivo*.

In order to increase the yield of magnetic AP recordings using our miniaturized GMR sensors, it would be very helpful to have electrodes integrated on the same probe as the GMR sensors. The two types of sensors would then be on the same needle and in one plane and could be brought very close together. The probes we used did contain electrode sites, but these were not functional due to technical difficulties during probe production. Future efforts might be directed towards integrating functional electrode sites without introducing additional noise in the GMR.

Alternatively, adding multiple additional magnetic sensors could also improve the yield of magnetic AP recordings. While this would increase the possibility to be in close vicinity to neurons and, using different sensor orientations, to record signals from neurons with arbitrary spatial orientations, this advancement would come with its own set of challenges. All GMR sensors would need to be connected with separate sets of contact lines, and the number of these lines that can be implemented within one probe is limited. Also, the more GMRs there are inside the tissue, the more substantial the sum of the small temperature increases caused by each individual sensor will be. While this is not of concern when using a small number of sensors (like in our current probe design), it already has an effect on spike rate and any further heating of the tissue has been shown, also in this work, to affect the spiking behavior of neurons in a substantial manner. Thus, while it would be beneficial to integrate more GMR sensors, it currently is not straightforward to implement.

Another major improvement would be an increase in the sensitivity of the GMR sensors themselves. In the existing probes, sensitivity can only be improved by increasing the current through the sensor. If this could be achieved without temperature increases, the sensor would be capable of detecting smaller fields than it currently can and thus reduce the need to average across a large number of spike events to detect the magnetic signature. This would probably facilitate the recording of magnetic AP signature more reliably and in a larger number of units. The development of sensors based on magnetic tunnel junctions (Julliere, 1975; Yuasa et al., 2004; Sharma et al., 2017) might add another technical approach worth of consideration for the Magnetrode, because they exhibit high sensitivity but require lower bias currents. However, TMR sensors exhibit a larger low-frequency noise than GMR sensors, and the limit of detection at 1 kHz is of the same order of magnitude (Fermon and Van de Voorde, 2016).

In conclusion, we demonstrated that it is possible, to use current state-of-the-art GMR sensors to detect the magnetic signature of action potentials of single neurons. This opens a wide field of future research questions, exploring the advantages that magnetic signals provide over electric recordings and the added information one can gain by combining the two. We also identified a number of possible technological improvements, which would make such magnetic recordings easier to achieve and to upscale.

## Acknowledgments

The authors would like to thank Claudia Kernberger for help with illustrations.

This work was supported by DFG (FR2557/6-1-NeuroTMR), EU (FP7-600730-Magnetrodes to P.F.), National Institutes of Health (1U54MH091657-WU-Minn-Consortium-HCP to P.F.).

## Author contributions

Conceptualization, P.F., M.P.-L., C.F., F.J.K., and P.J.; Methodology: C.C., A.S., M.P.-L., and C.F.;

Investigation: F.J.K., P.J., C.C., A.S., M.P.-L., and C.F.; Analysis: F.J.K., P.J., M.P.-D., C.F., M.P.-L., and P.F.;

Writing – Original draft: F.J.K., M.P.-L., and P.F.; Writing – Review & Editing: All authors; Funding Acquisition: M.P.-L., and P.F.

